## Supplementary_materials for "Inability to switch from ARID1A-BAF to ARID1B-BAF impairs exit from pluripotency and commitment towards neural crest formation in *ARID1B*-related neurodevelopmental disorders": Full Membranes.pptx

### Slide 1
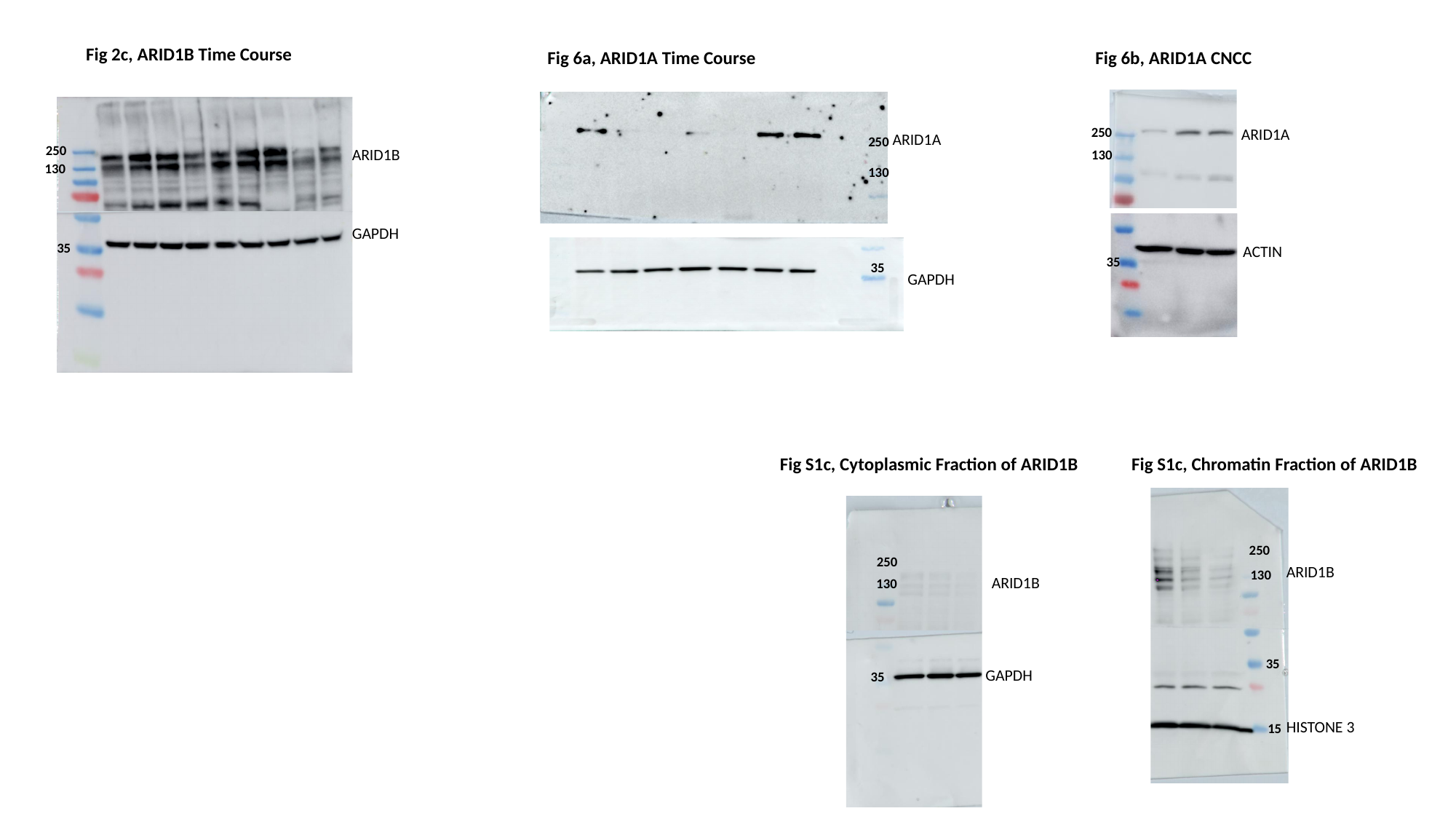

Fig 2c, ARID1B Time Course
Fig 6b, ARID1A CNCC
Fig 6a, ARID1A Time Course
250
ARID1A
ARID1A
250
250
ARID1B
130
130
130
GAPDH
35
ACTIN
35
35
GAPDH
Fig S1c, Cytoplasmic Fraction of ARID1B
Fig S1c, Chromatin Fraction of ARID1B
250
250
ARID1B
130
ARID1B
130
35
GAPDH
35
HISTONE 3
15
