## Supplementary_materials for "Inability to switch from ARID1A-BAF to ARID1B-BAF impairs exit from pluripotency and commitment towards neural crest formation in *ARID1B*-related neurodevelopmental disorders": Supplementary_File_S2_MOTIF_ANALYSIS_PATIENT_SPECIFIC_ATAC_SEQ_PEAKS.html

MEME ChIP


The name of the alphabet symbol.

[close ]

The frequency of the alphabet symbol as defined by the background model.

[close ]

MEME-ChIP outputs a tab-separated values (TSV) file ('summary.tsv') that
contains one line for each motif found by MEME-ChIP.
The lines are sorted in order of decreasing statistical significance.
The first line in the file contains the (tab-separated) names of the fields.
Your command line is given at the end of the file in a comment line starting with the
character '#'.
The names and meanings of the fields in the are described in the table below.

| field | name | contents |
| --- | --- | --- |
| 1 | MOTIF\_INDEX | The index of the motif in the "Motifs in MEME text format" file ('combined.meme') output by MEME-ChIP. |
| 2 | MOTIF\_SOURCE | The name of the program that found the *de novo* motif, or the name of the motif file containing the known motif. |
| 3 | MOTIF\_ID | The name of the motif, which is unique in the motif database file. |
| 4 | ALT\_ID | An alternate name for the motif, which may be provided in the motif database file. |
| 5 | CONSENSUS | The ID of the *de novo* motif, or a consensus sequence computed from the letter frequencies in the known motif (as described below). |
| 6 | WIDTH | The width of the motif. |
| 7 | SITES | The number of sites reported by the *de novo* program, or the number of "Total Matches" reported by CentriMo. |
| 8 | E-VALUE | The statistical significance of the motif. |
| 9 | E-VALUE\_SOURCE | The program that reported the *E*-value. |
| 10 | MOST\_SIMILAR\_MOTIF | The known motif most similar to this motif according to Tomtom. |
| 11 | URL | A link to a description of the most similar motif, or to the known motif. |

A **consensus sequence** is constructed from each column in a
motif's frequency matrix using the **"50% rule"**
as follows:

1. The letter frequencies in the column are sorted in decreasing order.
2. Letters with frequency less 50% of the maximum are discarded.
3. The letter used in this position in the consensus sequence is determined
   by the first rule below that applies:

- If there is only one letter left, or if the remaining letters exactly match
  an ambiguous symbol in the alphabet, the **letter** or **ambiguous symbol**,
  respectively, is used.
- Otherwise, if the remaining set contains at least 50% of the core
  symbols in the alphabet, the alphabet's **wildcard**
  (e.g., "N" for DNA or RNA, and "X" for protein) is used.
- Otherwise, the letter with the **maximum frequency** is used.

[
close ]

MEME-ChIP outputs a text file ('combined.meme') containing all the significant motifs found by MEME-ChIP.
The motifs are in Minimal MEME Motif format,
and their IDs correspond to the motif indices given in the "Summary in TSV Format" file ('summary.tsv').

**Note:** The 'nsites=' and 'E=' fields in the motif headers are only
relevant for the MEME and DREME motifs. For known motifs, those values do
not refer to the number of sites in the input sequences.

[
close ]

This is a link to the motif in the output of the particular motif
discovery (e.g., MEME) or motif enrichment (e.g., CentriMo) program that
reported it.

[
close ]

This is the significance of the motif according to the particular motif
discovery (e.g., MEME) or motif enrichment (e.g., CentriMo) program that
reported it.

Follow the link under the "Discovery/Enrichment Program" column for
more information on how the significance value was derived.

[
close ]

Motifs reported by a motif discovery program (e.g., MEME) are compared
with known motifs in a motif database specified by the user. This column
lists the (up to) three most similar motifs. Only known motifs with
TOMTOM similarity E-values of less than 1.0 to the discovered motif will
be shown here. Clicking any of these links will show the TOMTOM results
where all alignments can be viewed.

Motifs reported by a motif enrichment program (e.g., CentriMo) list the
motif's name and a link to the motif's entry on the database website if it
is available.

[
close ]

This graph shows the distribution of the best matches to the motif in
the sequences as found by a CentriMo analysis.

The vertical line in the center of the graph corresponds to the center
of the sequences.

Clicking on a motif's graph will take you to the CentriMo output with
that motif selected for graphing.

[
close ]

Clicking here will show you all the motifs found by motif discovery or
motif enrichment analysis that are significantly similar to the reported
motif.

The additional motifs are shown aligned with the reported motif,
sorted in order of significance of the motif according to the
particular motif discovery (e.g., MEME) or motif enrichment
(e.g., CentriMo) program that reported it.

To cluster the motifs MEME ChIP does the following:

1. Start with no groups and all significant reported motifs.
2. Run TOMTOM with all significant reported motifs to determine
   pairwise similarity.
3. Group Highly Similar Motifs---

   While ungrouped motifs:

   Select most significant ungrouped motif.

   This is called the "seed" motif for the group and we will call the
   E-value of its seed motif the group's "significance".

   Form a new group from the seed motif and all other motifs that
   are not yet in a group and who are strongly similar to the seed
   motif (default: TOMTOM E-value ≤ 0.05).
4. Merge Groups---

   For each group (most significant to least significant), merge it with
   any less significant group if all of its motifs are weakly similar to
   the first group's seed motif (default: TOMTOM E-value ≤ 0.1).

[
close ]

Clicking here takes you to the CentriMo motif enrichment analysis with
the results for this all the motifs in this group.

[
close ]

This lists links to related content, which may include:

- Motif Spacing Analysis--SpaMo results using this motif as
  the "primary" motif, and each of the discovered motifs and
  motifs in any motif databases specified to MEME-ChIP as
  potential "secondary" motifs. SpaMo reports the secondary
  motifs whose occurrences are enriched at particular distances relative
  to the primary motif's occurrences in the input sequences.
- Motif Sites in GFF3--FIMO results showing the positions of occurrences
  of this motif in the input sequences in GFF3
  format. If the input sequences to MEME-ChIP have FASTA headers following
  the UCSC style ("chromosome\_name:starting\_position-ending\_position"),
  and the chromosome names are in UCSC (not ENSEMBL) format,
  the GFF3 output will be suitable for uploading to the UCSC Genome Browser
  as a custom track.

[
close ]

### MEME-ChIP

#### Motif Analysis of Large Nucleotide Datasets

For further information on how to interpret these results please access
http://meme-suite.org//doc/meme-chip-output-format.html.  
To get a copy of the MEME software please access
http://meme-suite.org.

If you use MEME-ChIP in your research, please cite the following paper:  

Philip Machanick and Timothy L. Bailey, "MEME-ChIP: motif analysis of large DNA datasets",
*Bioinformatics*, **27**12, 1696-1697, 2011.
[full text]

Motifs
  |  
Programs
  |  
Input Files
  |  
Program information
  |  
Summary in TSV Format 

 
  |  
Motifs in MEME Text Format


### Javascript is required to view these results!

### Your browser does not support canvas!


#### Description

patient specific long


#### Motifs

**The significant motifs
(E-value ≤ 0.05)
found by the programs MEME, DREME and CentriMo;
clustered by similarity and ordered by E-value.**

Expand All Clusters
Collapse All Clusters

| Motif Found | Discovery/​Enrichment Program | E-value | Known or Similar Motifs | Distribution | SpaMo & FIMO |
| --- | --- | --- | --- | --- | --- |
|  | DREME | 6.0e-103 | CTCF\_HUMAN.H11MO.0.A INSM1\_HUMAN.H11MO.0.C CTCFL\_HUMAN.H11MO.0.A | Not Centrally Enriched | - Motif Spacing Analysis - Motif Sites in GFF3 |
|  | MEME | 3.8e-056 | CTCF\_HUMAN.H11MO.0.A CTCFL\_HUMAN.H11MO.0.A ZIC3\_HUMAN.H11MO.0.B | Not Centrally Enriched |  |

Reverse Complement ⇆Show 1 More ↧Show Less ↥

| Motif Found | Discovery/​Enrichment Program | E-value | Known or Similar Motifs | Distribution | SpaMo & FIMO |
| --- | --- | --- | --- | --- | --- |
|  | DREME | 1.9e-091 | SOX2\_HUMAN.H11MO.0.A SOX3\_HUMAN.H11MO.0.B SOX9\_HUMAN.H11MO.0.B | Not Centrally Enriched | - Motif Spacing Analysis - Motif Sites in GFF3 |

Reverse Complement ⇆

| Motif Found | Discovery/​Enrichment Program | E-value | Known or Similar Motifs | Distribution | SpaMo & FIMO |
| --- | --- | --- | --- | --- | --- |
|  | DREME | 2.9e-043 | NANOG\_HUMAN.H11MO.0.A PO5F1\_HUMAN.H11MO.0.A PO2F2\_HUMAN.H11MO.0.A | Not Centrally Enriched | - Motif Spacing Analysis - Motif Sites in GFF3 |
|  | DREME | 7.1e-006 | PO5F1\_HUMAN.H11MO.0.A NANOG\_HUMAN.H11MO.0.A PO2F2\_HUMAN.H11MO.0.A | Not Centrally Enriched |  |

Reverse Complement ⇆Show 1 More ↧Show Less ↥

| Motif Found | Discovery/​Enrichment Program | E-value | Known or Similar Motifs | Distribution | SpaMo & FIMO |
| --- | --- | --- | --- | --- | --- |
|  | DREME | 2.7e-039 | VEZF1\_HUMAN.H11MO.0.C ZN436\_HUMAN.H11MO.0.C IKZF1\_HUMAN.H11MO.0.C | Not Centrally Enriched | - Motif Spacing Analysis - Motif Sites in GFF3 |

Reverse Complement ⇆

| Motif Found | Discovery/​Enrichment Program | E-value | Known or Similar Motifs | Distribution | SpaMo & FIMO |
| --- | --- | --- | --- | --- | --- |
|  | DREME | 7.7e-021 | PRGR\_HUMAN.H11MO.0.A | Not Centrally Enriched | - Motif Spacing Analysis - Motif Sites in GFF3 |

Reverse Complement ⇆

| Motif Found | Discovery/​Enrichment Program | E-value | Known or Similar Motifs | Distribution | SpaMo & FIMO |
| --- | --- | --- | --- | --- | --- |
|  | DREME | 4.1e-016 | ZSC31\_HUMAN.H11MO.0.C ZIC3\_HUMAN.H11MO.0.B OSR2\_HUMAN.H11MO.0.C | Not Centrally Enriched | - Motif Spacing Analysis - Motif Sites in GFF3 |

Reverse Complement ⇆

| Motif Found | Discovery/​Enrichment Program | E-value | Known or Similar Motifs | Distribution | SpaMo & FIMO |
| --- | --- | --- | --- | --- | --- |
|  | DREME | 2.5e-015 | CTCF\_HUMAN.H11MO.0.A | Not Centrally Enriched | - Motif Spacing Analysis - Motif Sites in GFF3 |

Reverse Complement ⇆

| Motif Found | Discovery/​Enrichment Program | E-value | Known or Similar Motifs | Distribution | SpaMo & FIMO |
| --- | --- | --- | --- | --- | --- |
|  | MEME | 1.2e-013 | PRDM6\_HUMAN.H11MO.0.C ANDR\_HUMAN.H11MO.0.A FOXO4\_HUMAN.H11MO.0.C | Not Centrally Enriched | - Motif Spacing Analysis - Motif Sites in GFF3 |
|  | DREME | 1.2e-006 | ZN274\_HUMAN.H11MO.0.A PRDM6\_HUMAN.H11MO.0.C NFAC1\_HUMAN.H11MO.0.B | Not Centrally Enriched |  |

Reverse Complement ⇆Show 1 More ↧Show Less ↥

| Motif Found | Discovery/​Enrichment Program | E-value | Known or Similar Motifs | Distribution | SpaMo & FIMO |
| --- | --- | --- | --- | --- | --- |
|  | DREME | 1.7e-012 | FOXO1\_HUMAN.H11MO.0.A FOXK1\_HUMAN.H11MO.0.A FOXO3\_HUMAN.H11MO.0.B | Not Centrally Enriched | - Motif Spacing Analysis - Motif Sites in GFF3 |

Reverse Complement ⇆

| Motif Found | Discovery/​Enrichment Program | E-value | Known or Similar Motifs | Distribution | SpaMo & FIMO |
| --- | --- | --- | --- | --- | --- |
|  | DREME | 4.4e-011 |  | Not Centrally Enriched | - Motif Spacing Analysis - Motif Sites in GFF3 |

Reverse Complement ⇆

| Motif Found | Discovery/​Enrichment Program | E-value | Known or Similar Motifs | Distribution | SpaMo & FIMO |
| --- | --- | --- | --- | --- | --- |
|  | DREME | 4.7e-009 | KLF3\_HUMAN.H11MO.0.B SP4\_HUMAN.H11MO.0.A KLF12\_HUMAN.H11MO.0.C | Not Centrally Enriched | - Motif Spacing Analysis - Motif Sites in GFF3 |

Reverse Complement ⇆

| Motif Found | Discovery/​Enrichment Program | E-value | Known or Similar Motifs | Distribution | SpaMo & FIMO |
| --- | --- | --- | --- | --- | --- |
|  | DREME | 9.0e-009 | DLX3\_HUMAN.H11MO.0.C LHX2\_HUMAN.H11MO.0.A NOBOX\_HUMAN.H11MO.0.C | Not Centrally Enriched | - Motif Spacing Analysis - Motif Sites in GFF3 |

Reverse Complement ⇆

| Motif Found | Discovery/​Enrichment Program | E-value | Known or Similar Motifs | Distribution | SpaMo & FIMO |
| --- | --- | --- | --- | --- | --- |
|  | DREME | 9.1e-007 |  | Not Centrally Enriched | - Motif Spacing Analysis - Motif Sites in GFF3 |

Reverse Complement ⇆

| Motif Found | Discovery/​Enrichment Program | E-value | Known or Similar Motifs | Distribution | SpaMo & FIMO |
| --- | --- | --- | --- | --- | --- |
|  | DREME | 6.1e-006 | TAF1\_HUMAN.H11MO.0.A | Not Centrally Enriched | - Motif Spacing Analysis - Motif Sites in GFF3 |

Reverse Complement ⇆

| Motif Found | Discovery/​Enrichment Program | E-value | Known or Similar Motifs | Distribution | SpaMo & FIMO |
| --- | --- | --- | --- | --- | --- |
|  | DREME | 2.5e-002 | ZN436\_HUMAN.H11MO.0.C ZN528\_HUMAN.H11MO.0.C COE1\_HUMAN.H11MO.0.A | Not Centrally Enriched | - Motif Spacing Analysis - Motif Sites in GFF3 |

Reverse Complement ⇆

| Motif Found | Discovery/​Enrichment Program | E-value | Known or Similar Motifs | Distribution | SpaMo & FIMO |
| --- | --- | --- | --- | --- | --- |
|  | DREME | 2.9e-002 | EVI1\_HUMAN.H11MO.0.B | Not Centrally Enriched | - Motif Spacing Analysis - Motif Sites in GFF3 |

Reverse Complement ⇆

| Motif Found | Discovery/​Enrichment Program | E-value | Known or Similar Motifs | Distribution | SpaMo & FIMO |
| --- | --- | --- | --- | --- | --- |
|  | DREME | 4.0e-002 | NFIA\_HUMAN.H11MO.0.C | Not Centrally Enriched | - Motif Spacing Analysis - Motif Sites in GFF3 |

Reverse Complement ⇆

#### Programs

| Command | Running Time | Status | Outputs |
| --- | --- | --- | --- |
| **getsize** ./ATAC\_CNCC5\_PATIENT\_SPECIFIC\_rep2\_LONG\_VERSION.fasta 1> $metrics | 0.05s | Success |  |
| **fasta-most** -min 50 < ./ATAC\_CNCC5\_PATIENT\_SPECIFIC\_rep2\_LONG\_VERSION.fasta 1> $metrics | 0.12s | Success |  |
| **fasta-center** -dna -len 100 < ./ATAC\_CNCC5\_PATIENT\_SPECIFIC\_rep2\_LONG\_VERSION.fasta 1> ./seqs-centered | 0.19s | Success | - seqs-centered |
| **fasta-shuffle-letters** ./seqs-centered ./seqs-shuffled -kmer 2 -tag -dinuc -dna -seed 1 | 0.08s | Success | - seqs-shuffled |
| **fasta-get-markov** -nostatus -nosummary -dna -m 1 ./ATAC\_CNCC5\_PATIENT\_SPECIFIC\_rep2\_LONG\_VERSION.fasta ./background | 0.03s | Success | - Background |
| **meme** ./seqs-centered -oc meme\_out -mod zoops -nmotifs 12 -minw 6 -maxw 10 -bfile ./background -dna -searchsize 100000 -time 5082 -revcomp -nostatus | 1h 1m 45.79s | Success | - MEME HTML - MEME text - MEME XML |
| **dreme** -verbosity 1 -oc dreme\_out -png -dna -p ./seqs-centered -n ./seqs-shuffled -t 5657 -e 0.05 | 18m 6.85s | Success | - DREME HTML - DREME text - DREME XML |
| **centrimo** -seqlen 73 -verbosity 1 -oc centrimo\_out -bfile ./background -score 5.0 -ethresh 10.0 ./ATAC\_CNCC5\_PATIENT\_SPECIFIC\_rep2\_LONG\_VERSION.fasta meme\_out/meme.xml dreme\_out/dreme.xml db/HUMAN/HOCOMOCOv11\_core\_HUMAN\_mono\_meme\_format.meme | 2.14s | Warnings | - CentriMo HTML - Site Counts |
| **tomtom** -verbosity 1 -oc meme\_tomtom\_out -min-overlap 5 -dist pearson -evalue -thresh 1 -no-ssc meme\_out/meme.xml db/HUMAN/HOCOMOCOv11\_core\_HUMAN\_mono\_meme\_format.meme | 4.58s | Success | - Tomtom HTML - Tomtom TSV - Tomtom XML |
| **tomtom** -verbosity 1 -oc dreme\_tomtom\_out -min-overlap 5 -dist pearson -evalue -thresh 1 -no-ssc dreme\_out/dreme.xml db/HUMAN/HOCOMOCOv11\_core\_HUMAN\_mono\_meme\_format.meme | 3.84s | Success | - Tomtom HTML - Tomtom TSV - Tomtom XML |
| **tomtom** -verbosity 1 -text -thresh 0.1 ./combined.meme ./combined.meme 1> ./motif\_alignment.txt | 0.14s | Success | - Motif Alignment |
| **spamo** -verbosity 1 -oc spamo\_out\_1 -bgfile ./background -keepprimary -primary AGRKGGCR ./ATAC\_CNCC5\_PATIENT\_SPECIFIC\_rep2\_LONG\_VERSION.fasta dreme\_out/dreme.xml meme\_out/meme.xml dreme\_out/dreme.xml db/HUMAN/HOCOMOCOv11\_core\_HUMAN\_mono\_meme\_format.meme | 3.50s | Warnings | - SpaMo HTML |
| **spamo** -verbosity 1 -oc spamo\_out\_2 -bgfile ./background -keepprimary -primary ACAAWRV ./ATAC\_CNCC5\_PATIENT\_SPECIFIC\_rep2\_LONG\_VERSION.fasta dreme\_out/dreme.xml meme\_out/meme.xml dreme\_out/dreme.xml db/HUMAN/HOCOMOCOv11\_core\_HUMAN\_mono\_meme\_format.meme | 5.14s | Warnings | - SpaMo HTML |
| **spamo** -verbosity 1 -oc spamo\_out\_3 -bgfile ./background -keepprimary -primary ATGYWAAT ./ATAC\_CNCC5\_PATIENT\_SPECIFIC\_rep2\_LONG\_VERSION.fasta dreme\_out/dreme.xml meme\_out/meme.xml dreme\_out/dreme.xml db/HUMAN/HOCOMOCOv11\_core\_HUMAN\_mono\_meme\_format.meme | 2.58s | Warnings | - SpaMo HTML |
| **spamo** -verbosity 1 -oc spamo\_out\_4 -bgfile ./background -keepprimary -primary RGGARR ./ATAC\_CNCC5\_PATIENT\_SPECIFIC\_rep2\_LONG\_VERSION.fasta dreme\_out/dreme.xml meme\_out/meme.xml dreme\_out/dreme.xml db/HUMAN/HOCOMOCOv11\_core\_HUMAN\_mono\_meme\_format.meme | 12.96s | Warnings | - SpaMo HTML |
| **spamo** -verbosity 1 -oc spamo\_out\_5 -bgfile ./background -keepprimary -primary RAATR ./ATAC\_CNCC5\_PATIENT\_SPECIFIC\_rep2\_LONG\_VERSION.fasta dreme\_out/dreme.xml meme\_out/meme.xml dreme\_out/dreme.xml db/HUMAN/HOCOMOCOv11\_core\_HUMAN\_mono\_meme\_format.meme | 9.41s | Warnings | - SpaMo HTML |
| **spamo** -verbosity 1 -oc spamo\_out\_6 -bgfile ./background -keepprimary -primary CHGCAG ./ATAC\_CNCC5\_PATIENT\_SPECIFIC\_rep2\_LONG\_VERSION.fasta dreme\_out/dreme.xml meme\_out/meme.xml dreme\_out/dreme.xml db/HUMAN/HOCOMOCOv11\_core\_HUMAN\_mono\_meme\_format.meme | 6.43s | Warnings | - SpaMo HTML |
| **spamo** -verbosity 1 -oc spamo\_out\_7 -bgfile ./background -keepprimary -primary CCACTAGR ./ATAC\_CNCC5\_PATIENT\_SPECIFIC\_rep2\_LONG\_VERSION.fasta dreme\_out/dreme.xml meme\_out/meme.xml dreme\_out/dreme.xml db/HUMAN/HOCOMOCOv11\_core\_HUMAN\_mono\_meme\_format.meme | 1.88s | Warnings | - SpaMo HTML |
| **spamo** -verbosity 1 -oc spamo\_out\_8 -bgfile ./background -keepprimary -primary TTTKTTTTTT ./ATAC\_CNCC5\_PATIENT\_SPECIFIC\_rep2\_LONG\_VERSION.fasta meme\_out/meme.xml meme\_out/meme.xml dreme\_out/dreme.xml db/HUMAN/HOCOMOCOv11\_core\_HUMAN\_mono\_meme\_format.meme | 4.86s | Warnings | - SpaMo HTML |
| **spamo** -verbosity 1 -oc spamo\_out\_9 -bgfile ./background -keepprimary -primary GDAAACA ./ATAC\_CNCC5\_PATIENT\_SPECIFIC\_rep2\_LONG\_VERSION.fasta dreme\_out/dreme.xml meme\_out/meme.xml dreme\_out/dreme.xml db/HUMAN/HOCOMOCOv11\_core\_HUMAN\_mono\_meme\_format.meme | 3.01s | Warnings | - SpaMo HTML |
| **spamo** -verbosity 1 -oc spamo\_out\_10 -bgfile ./background -keepprimary -primary CTGKGW ./ATAC\_CNCC5\_PATIENT\_SPECIFIC\_rep2\_LONG\_VERSION.fasta dreme\_out/dreme.xml meme\_out/meme.xml dreme\_out/dreme.xml db/HUMAN/HOCOMOCOv11\_core\_HUMAN\_mono\_meme\_format.meme | 6.34s | Warnings | - SpaMo HTML |
| **spamo** -verbosity 1 -oc spamo\_out\_11 -bgfile ./background -keepprimary -primary GCYCCRCC ./ATAC\_CNCC5\_PATIENT\_SPECIFIC\_rep2\_LONG\_VERSION.fasta dreme\_out/dreme.xml meme\_out/meme.xml dreme\_out/dreme.xml db/HUMAN/HOCOMOCOv11\_core\_HUMAN\_mono\_meme\_format.meme | 3.44s | Warnings | - SpaMo HTML |
| **spamo** -verbosity 1 -oc spamo\_out\_12 -bgfile ./background -keepprimary -primary STAATTA ./ATAC\_CNCC5\_PATIENT\_SPECIFIC\_rep2\_LONG\_VERSION.fasta dreme\_out/dreme.xml meme\_out/meme.xml dreme\_out/dreme.xml db/HUMAN/HOCOMOCOv11\_core\_HUMAN\_mono\_meme\_format.meme | 2.66s | Warnings | - SpaMo HTML |
| **spamo** -verbosity 1 -oc spamo\_out\_13 -bgfile ./background -keepprimary -primary GYGGTR ./ATAC\_CNCC5\_PATIENT\_SPECIFIC\_rep2\_LONG\_VERSION.fasta dreme\_out/dreme.xml meme\_out/meme.xml dreme\_out/dreme.xml db/HUMAN/HOCOMOCOv11\_core\_HUMAN\_mono\_meme\_format.meme | 4.25s | Warnings | - SpaMo HTML |
| **spamo** -verbosity 1 -oc spamo\_out\_14 -bgfile ./background -keepprimary -primary CGCYGCCG ./ATAC\_CNCC5\_PATIENT\_SPECIFIC\_rep2\_LONG\_VERSION.fasta dreme\_out/dreme.xml meme\_out/meme.xml dreme\_out/dreme.xml db/HUMAN/HOCOMOCOv11\_core\_HUMAN\_mono\_meme\_format.meme | 2.65s | Warnings | - SpaMo HTML |
| **spamo** -verbosity 1 -oc spamo\_out\_15 -bgfile ./background -keepprimary -primary CMGGGA ./ATAC\_CNCC5\_PATIENT\_SPECIFIC\_rep2\_LONG\_VERSION.fasta dreme\_out/dreme.xml meme\_out/meme.xml dreme\_out/dreme.xml db/HUMAN/HOCOMOCOv11\_core\_HUMAN\_mono\_meme\_format.meme | 5.97s | Warnings | - SpaMo HTML |
| **spamo** -verbosity 1 -oc spamo\_out\_16 -bgfile ./background -keepprimary -primary AGAYAAT ./ATAC\_CNCC5\_PATIENT\_SPECIFIC\_rep2\_LONG\_VERSION.fasta dreme\_out/dreme.xml meme\_out/meme.xml dreme\_out/dreme.xml db/HUMAN/HOCOMOCOv11\_core\_HUMAN\_mono\_meme\_format.meme | 2.53s | Warnings | - SpaMo HTML |
| **spamo** -verbosity 1 -oc spamo\_out\_17 -bgfile ./background -keepprimary -primary CACGGAGY ./ATAC\_CNCC5\_PATIENT\_SPECIFIC\_rep2\_LONG\_VERSION.fasta dreme\_out/dreme.xml meme\_out/meme.xml dreme\_out/dreme.xml db/HUMAN/HOCOMOCOv11\_core\_HUMAN\_mono\_meme\_format.meme | 1.80s | Warnings | - SpaMo HTML |
| **fimo** --parse-genomic-coord --verbosity 1 --oc fimo\_out\_1 --bgfile ./background --motif AGRKGGCR dreme\_out/dreme.xml ./ATAC\_CNCC5\_PATIENT\_SPECIFIC\_rep2\_LONG\_VERSION.fasta | 0.83s | Success | - FIMO GFF - FIMO HTML - FIMO TSV |
| **fimo** --parse-genomic-coord --verbosity 1 --oc fimo\_out\_2 --bgfile ./background --motif ACAAWRV dreme\_out/dreme.xml ./ATAC\_CNCC5\_PATIENT\_SPECIFIC\_rep2\_LONG\_VERSION.fasta | 0.83s | Success | - FIMO GFF - FIMO HTML - FIMO TSV |
| **fimo** --parse-genomic-coord --verbosity 1 --oc fimo\_out\_3 --bgfile ./background --motif ATGYWAAT dreme\_out/dreme.xml ./ATAC\_CNCC5\_PATIENT\_SPECIFIC\_rep2\_LONG\_VERSION.fasta | 0.83s | Success | - FIMO GFF - FIMO HTML - FIMO TSV |
| **fimo** --parse-genomic-coord --verbosity 1 --oc fimo\_out\_4 --bgfile ./background --motif RGGARR dreme\_out/dreme.xml ./ATAC\_CNCC5\_PATIENT\_SPECIFIC\_rep2\_LONG\_VERSION.fasta | 0.83s | Success | - FIMO GFF - FIMO HTML - FIMO TSV |
| **fimo** --parse-genomic-coord --verbosity 1 --oc fimo\_out\_5 --bgfile ./background --motif RAATR dreme\_out/dreme.xml ./ATAC\_CNCC5\_PATIENT\_SPECIFIC\_rep2\_LONG\_VERSION.fasta | 0.78s | Success | - FIMO GFF - FIMO HTML - FIMO TSV |
| **fimo** --parse-genomic-coord --verbosity 1 --oc fimo\_out\_6 --bgfile ./background --motif CHGCAG dreme\_out/dreme.xml ./ATAC\_CNCC5\_PATIENT\_SPECIFIC\_rep2\_LONG\_VERSION.fasta | 0.79s | Success | - FIMO GFF - FIMO HTML - FIMO TSV |
| **fimo** --parse-genomic-coord --verbosity 1 --oc fimo\_out\_7 --bgfile ./background --motif CCACTAGR dreme\_out/dreme.xml ./ATAC\_CNCC5\_PATIENT\_SPECIFIC\_rep2\_LONG\_VERSION.fasta | 0.82s | Success | - FIMO GFF - FIMO HTML - FIMO TSV |
| **fimo** --parse-genomic-coord --verbosity 1 --oc fimo\_out\_8 --bgfile ./background --motif TTTKTTTTTT meme\_out/meme.xml ./ATAC\_CNCC5\_PATIENT\_SPECIFIC\_rep2\_LONG\_VERSION.fasta | 0.84s | Success | - FIMO GFF - FIMO HTML - FIMO TSV |
| **fimo** --parse-genomic-coord --verbosity 1 --oc fimo\_out\_9 --bgfile ./background --motif GDAAACA dreme\_out/dreme.xml ./ATAC\_CNCC5\_PATIENT\_SPECIFIC\_rep2\_LONG\_VERSION.fasta | 0.82s | Success | - FIMO GFF - FIMO HTML - FIMO TSV |
| **fimo** --parse-genomic-coord --verbosity 1 --oc fimo\_out\_10 --bgfile ./background --motif CTGKGW dreme\_out/dreme.xml ./ATAC\_CNCC5\_PATIENT\_SPECIFIC\_rep2\_LONG\_VERSION.fasta | 0.79s | Success | - FIMO GFF - FIMO HTML - FIMO TSV |
| **fimo** --parse-genomic-coord --verbosity 1 --oc fimo\_out\_11 --bgfile ./background --motif GCYCCRCC dreme\_out/dreme.xml ./ATAC\_CNCC5\_PATIENT\_SPECIFIC\_rep2\_LONG\_VERSION.fasta | 0.82s | Success | - FIMO GFF - FIMO HTML - FIMO TSV |
| **fimo** --parse-genomic-coord --verbosity 1 --oc fimo\_out\_12 --bgfile ./background --motif STAATTA dreme\_out/dreme.xml ./ATAC\_CNCC5\_PATIENT\_SPECIFIC\_rep2\_LONG\_VERSION.fasta | 0.82s | Success | - FIMO GFF - FIMO HTML - FIMO TSV |
| **fimo** --parse-genomic-coord --verbosity 1 --oc fimo\_out\_13 --bgfile ./background --motif GYGGTR dreme\_out/dreme.xml ./ATAC\_CNCC5\_PATIENT\_SPECIFIC\_rep2\_LONG\_VERSION.fasta | 0.79s | Success | - FIMO GFF - FIMO HTML - FIMO TSV |
| **fimo** --parse-genomic-coord --verbosity 1 --oc fimo\_out\_14 --bgfile ./background --motif CGCYGCCG dreme\_out/dreme.xml ./ATAC\_CNCC5\_PATIENT\_SPECIFIC\_rep2\_LONG\_VERSION.fasta | 0.81s | Success | - FIMO GFF - FIMO HTML - FIMO TSV |
| **fimo** --parse-genomic-coord --verbosity 1 --oc fimo\_out\_15 --bgfile ./background --motif CMGGGA dreme\_out/dreme.xml ./ATAC\_CNCC5\_PATIENT\_SPECIFIC\_rep2\_LONG\_VERSION.fasta | 0.79s | Success | - FIMO GFF - FIMO HTML - FIMO TSV |
| **fimo** --parse-genomic-coord --verbosity 1 --oc fimo\_out\_16 --bgfile ./background --motif AGAYAAT dreme\_out/dreme.xml ./ATAC\_CNCC5\_PATIENT\_SPECIFIC\_rep2\_LONG\_VERSION.fasta | 0.81s | Success | - FIMO GFF - FIMO HTML - FIMO TSV |
| **fimo** --parse-genomic-coord --verbosity 1 --oc fimo\_out\_17 --bgfile ./background --motif CACGGAGY dreme\_out/dreme.xml ./ATAC\_CNCC5\_PATIENT\_SPECIFIC\_rep2\_LONG\_VERSION.fasta | 1.08s | Success | - FIMO GFF - FIMO HTML - FIMO TSV |

#### Input Files

###### Alphabet

    **Background source:** built from the (primary) sequences

| Name | Bg. |  |  |  | Bg. | Name |
| --- | --- | --- | --- | --- | --- | --- |
| Adenine | 0.2304 | A | ~ | T | 0.2304 | Thymine |
| Cytosine | 0.2696 | C | ~ | G | 0.2696 | Guanine |

###### Primary Sequences

| Database | Source | Sequence Count |
| --- | --- | --- |
| ATAC CNCC5 PATIENT SPECIFIC rep2 LONG VERSION | ATAC\_CNCC5\_PATIENT\_SPECIFIC\_rep2\_LONG\_VERSION.fasta | 8144 |

###### Motifs

| Database | Source | Motif Count |
| --- | --- | --- |
| HOCOMOCOv11 core HUMAN mono meme format | db/HUMAN/HOCOMOCOv11\_core\_HUMAN\_mono\_meme\_format.meme | 401 |

###### MEME-ChIP version

5.2.0
(Release date: Wed Oct 14 12:02:54 2020 -0700)

###### Command line summary
