## Supplementary_materials for "Inability to switch from ARID1A-BAF to ARID1B-BAF impairs exit from pluripotency and commitment towards neural crest formation in *ARID1B*-related neurodevelopmental disorders": Supplementary_File_S2_MOTIF_ANALYSIS_PATIENT_SPECIFIC_ATAC_SEQ_PEAKS.PDF

### MEME-ChIP
Motif Analysis of Large Nucleotide Datasets

For further information on how to interpret these results please access [http://meme-suite.org//doc/meme-chip-output-format.html](http://meme-suite.org//doc/meme-chip-output-format.html).
To get a copy of the MEME software please access [http://meme-suite.org](http://meme-suite.org).

If you use MEME-ChIP in your research, please cite the following paper:
Philip Machanick and Timothy L. Bailey, "MEME-ChIP: motif analysis of large DNA datasets", *Bioinformatics*, **27**12, 1696-1697, 2011. [full text]

MOTIFS | PROGRAMS | INPUT FILES | PROGRAM INFORMATION | SUMMARY IN TSV FORMAT NEW | MOTIFS IN MEME TEXT FORMAT NEW

#### DESCRIPTION

patient specific long

#### MOTIFS

**The significant motifs (E-value ≤ 0.05) found by the programs MEME, DREME and CentriMo; clustered by similarity and ordered by E-value.**

Expand All Clusters Collapse All Clusters

| Motif Found | Discovery/ Program | E-value | Known or Similar Motifs | Distribution | SpaMo FIMO |
|---|---|---|---|---|---|
| 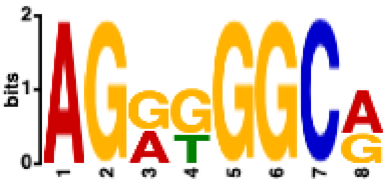 | DREME | 6.0e-103 | CTCF_HUMAN.H11MO.0.A<br>INSM1_HUMAN.H11MO.0.C<br>CTCFL_HUMAN.H11MO.0.A | Not Centrally Enriched | • Mot Spa Ana<br>• Mot Site GFF |

Reverse Complement ⇆ Show 1 More ↧

| Motif Found | Discovery/ Program | E-value | Known or Similar Motifs | Distribution | SpaMo & FIMO |
|---|---|---|---|---|---|
| 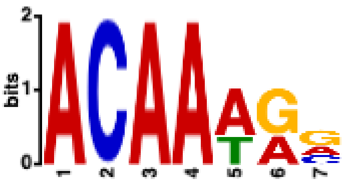 | DREME | 1.9e-091 | SOX2_HUMAN.H11MO.0.A<br>SOX3_HUMAN.H11MO.0.B<br>SOX9_HUMAN.H11MO.0.B | Not Centrally Enriched | • Motif Spacing Analysis<br>• Motif Sites in GFF3 |

Reverse Complement ⇆

| Motif Found | Discovery/ Enrichment Program | E-value | Known or Similar Motifs | Distribution | SpaMo FIMO |
|---|---|---|---|---|---|
| 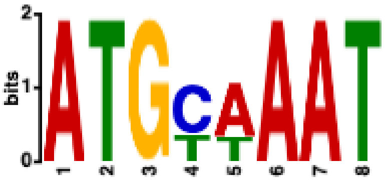 | DREME | 2.9e-043 | NANOG_HUMAN.H11MO.0.A<br>PO5F1_HUMAN.H11MO.0.A<br>PO2F2_HUMAN.H11MO.0.A | Not Centrally Enriched | • Mo Spa Ana<br>• Mo Site GFF |

Reverse Complement ⇆ Show 1 More ↧

| Motif Found | Discovery/ Enrichment Program | E-value | Known or Similar Motifs | Distribution | SpaMo & FIMO |
|---|---|---|---|---|---|
| 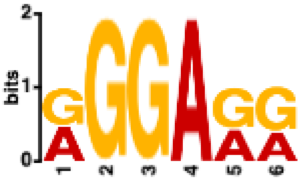 | DREME | 2.7e-039 | VEZF1_HUMAN.H11MO.0.C<br>ZN436_HUMAN.H11MO.0.C<br>IKZF1_HUMAN.H11MO.0.C | Not Centrally Enriched | • Motif Spacing Analysis<br>• Motif Sites in GFF3 |

Reverse Complement ⇆

| Motif Found | Discovery/ Enrichment Program | E-value | Known or Similar Motifs | Distribution | SpaMo & FIMO |
|---|---|---|---|---|---|
| 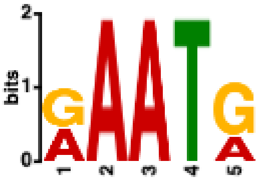 | DREME | 7.7e-021 | PRGR_HUMAN.H11MO.0.A | Not Centrally Enriched | • Motif Spacing Analysis<br>• Motif Sites in GFF3 |

Reverse Complement ⇆

| Motif Found | Discovery/ Enrichment Program | E-value | Known or Similar Motifs | Distribution | SpaMo & FIMO |
|---|---|---|---|---|---|
| 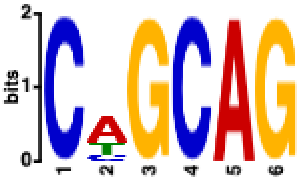 | DREME | 4.1e-016 | ZSC31_HUMAN.H11MO.0.C<br>ZIC3_HUMAN.H11MO.0.B<br>OSR2_HUMAN.H11MO.0.C | Not Centrally Enriched | • Motif Spacing Analysis<br>• Motif Sites in GFF3 |

Reverse Complement ⇆

| Motif Found | Discovery/ Enrichment Program | E-value | Known or Similar Motifs | Distribution | SpaMo & FIMO |
|---|---|---|---|---|---|

| Motif Found | Discovery/ Enrichment Program | E-value | Known or Similar Motifs | Distribution | SpaMo & FIMO |
|---|---|---|---|---|---|
| 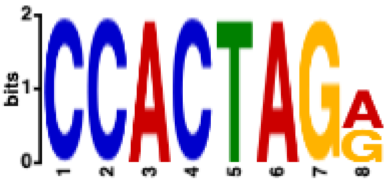 | DREME | 2.5e-015 | CTCF_HUMAN.H11MO.0.A | Not Centrally Enriched | • Motif Spaci Analy<br>• Motif Sites GFF3 |

Reverse Complement ⇆

| Motif Found | Discovery/ Enrichment Program | E-value | Known or Similar Motifs | Distribution | S |
|---|---|---|---|---|---|
| 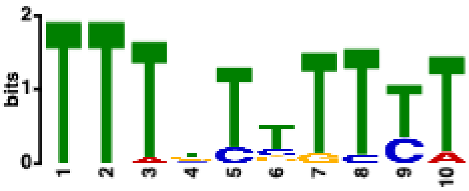 | MEME | 1.2e-013 | PRDM6_HUMAN.H11MO.0.C<br>ANDR_HUMAN.H11MO.0.A<br>FOXO4_HUMAN.H11MO.0.C | Not Centrally Enriched | |

Reverse Complement ⇆ Show 1 More ↧

| Motif Found | Discovery/ Enrichment Program | E-value | Known or Similar Motifs | Distribution | SpaMo & FIMO |
|---|---|---|---|---|---|
| 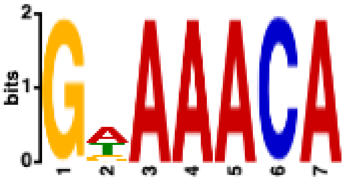 | DREME | 1.7e-012 | FOXO1_HUMAN.H11MO.0.A<br>FOXK1_HUMAN.H11MO.0.A<br>FOXO3_HUMAN.H11MO.0.B | Not Centrally Enriched | • Motif Spacing Analysi<br>• Motif Sites in GFF3 |

Reverse Complement ⇆

| Motif Found | Discovery/ Enrichment Program | E-value | Known or Similar Motifs | Distribution | SpaMo & FIMO |
|---|---|---|---|---|---|
| 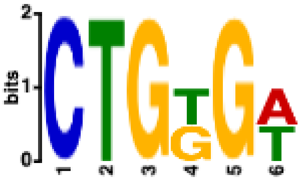 | DREME | 4.4e-011 | | Not Centrally Enriched | • Motif Spacing Analysis<br>• Motif Sites in GFF3 |

Reverse Complement ⇆

| Motif Found | Discovery/ Enrichment Program | E-value | Known or Similar Motifs | Distribution | SpaMo & FIMO |
|---|---|---|---|---|---|

| Motif Found | Discovery/ Enrichment Program | E-value | Known or Similar Motifs | Distribution | SpaMo & FIMO |
|---|---|---|---|---|---|
| 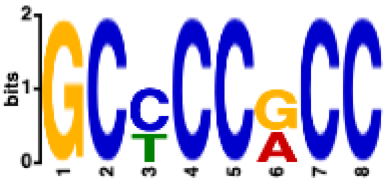 | DREME | 4.7e-009 | KLF3_HUMAN.H11MO.0.B<br>SP4_HUMAN.H11MO.0.A<br>KLF12_HUMAN.H11MO.0.C | Not Centrally Enriched | • Moti Spac Anal<br>• Moti Sites GFF3 |

Reverse Complement ⇆

| Motif Found | Discovery/ Enrichment Program | E-value | Known or Similar Motifs | Distribution | SpaMo & FIMO |
|---|---|---|---|---|---|
| 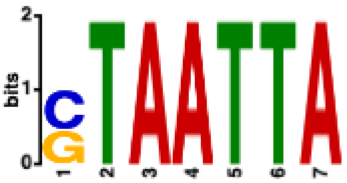 | DREME | 9.0e-009 | DLX3_HUMAN.H11MO.0.C<br>LHX2_HUMAN.H11MO.0.A<br>NOBOX_HUMAN.H11MO.0.C | Not Centrally Enriched | • Motif Spacin Analys<br>• Motif Sites ir GFF3 |

Reverse Complement ⇆

| Motif Found | Discovery/ Enrichment Program | E-value | Known or Similar Motifs | Distribution | SpaMo & FIMO |
|---|---|---|---|---|---|
| 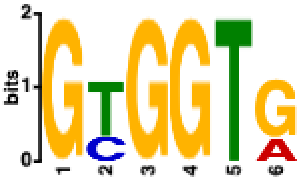 | DREME | 9.1e-007 | | Not Centrally Enriched | • Motif Spacing Analysis<br>• Motif Sites in GFF3 |

Reverse Complement ⇆

| Motif Found | Discovery/ Enrichment Program | E-value | Known or Similar Motifs | Distribution | SpaMo & FIMO |
|---|---|---|---|---|---|
| 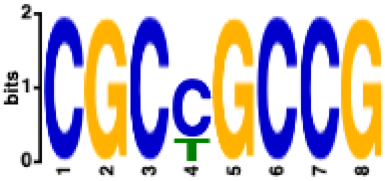 | DREME | 6.1e-006 | TAF1_HUMAN.H11MO.0.A | Not Centrally Enriched | • Motif Spaci Analy<br>• Motif Sites GFF3 |

Reverse Complement ⇆

| Motif Found | Discovery/ Enrichment Program | E-value | Known or Similar Motifs | Distribution | SpaMo & FIMO |
|---|---|---|---|---|---|

| Motif Found | Discovery/ Program | E-Enrichment value | Known or Similar Motifs | Distribution | SpaMo & FIMO |
|---|---|---|---|---|---|
| 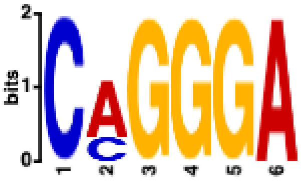 | DREME | 2.5e-002 | ZN436_HUMAN.H11MO.0.C<br>ZN528_HUMAN.H11MO.0.C<br>COE1_HUMAN.H11MO.0.A | Not Centrally Enriched | • Motif Spacing Analysis<br>• Motif Sites in GFF3 |

Reverse Complement ⇆

| Motif Found | Discovery/ Program | E-Enrichment value | Known or Similar Motifs | Distribution | SpaMo & FIMO |
|---|---|---|---|---|---|
| 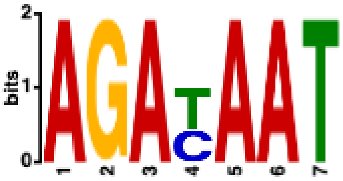 | DREME | 2.9e-002 | EVI1_HUMAN.H11MO.0.B | Not Centrally Enriched | • Motif Spacing Analysis<br>• Motif Sites in GFF3 |

Reverse Complement ⇆

| Motif Found | Discovery/ Program | E-Enrichment value | Known or Similar Motifs | Distribution | SpaMo & FIMO |
|---|---|---|---|---|---|
| 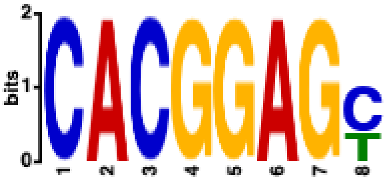 | DREME | 4.0e-002 | NFIA_HUMAN.H11MO.0.C | Not Centrally Enriched | • Motif Spaci Analy<br>• Motif Sites GFF3 |

```
meme-chip -oc . -time 300 -ccut 100 -fdesc description -order 1 -db
db/HUMAN/HOCOMOCOv11_core_HUMAN_mono_meme_format.meme -meme-mod zoops -meme-minw 6 -
meme-maxw 10 -meme-nmotifs 12 -meme-searchsize 100000 -dreme-e 0.05 -centrimo-score 5.0
-centrimo-ethresh 10.0 ATAC_CNCC5_PATIENT_SPECIFIC_rep2_LONG_VERSION.fasta
```
