## Supplementary_materials for "Inability to switch from ARID1A-BAF to ARID1B-BAF impairs exit from pluripotency and commitment towards neural crest formation in *ARID1B*-related neurodevelopmental disorders": Supplementary_Table_S1.docx

| **Supplementary Table S1 -LIST OF PRIMERS USED IN THIS STUDY** | | | |
| --- | --- | --- | --- |
| **TARGET** | **PRIMER NAME** | **SEQUENCE** | **APPLICATION** |
| 18s RNA | 18S forward | 5’-ATACATGCCGACGGGCGCTG-3’ | qRT-PCR |
|  | 18S reverse | 5’-AGGGGCTGACCGGGTTGGTT-3’ | qRT-PCR |
| SOX2 | SOX2 forward | 5’-GCCGAGTGGAAACTTTTGTCG-3’ | qRT-PCR |
|  | SOX2 reverse | 5’-GCAGCGTGTACTTATCCTTCTT-3’ | qRT-PCR |
| OCT4 | OCT4 forward | 5’-TCGAGAACCGAGTGAGAGG-3’ | qRT-PCR |
|  | OCT4 reverse | 5’-GAACCACACTCGGACCACA-3’ | qRT-PCR |
| NANOG | NANOG forward | 5’-ATGCCTCACACGGAGACTGT-3’ | qRT-PCR |
|  | NANOG reverse | 5’-AAGTGGGTTGTTTGCCTTTG-3’ | qRT-PCR |
| TFAP2A | TFAP2A forward | 5’-AACATGCTCCTGGCTACAAAA-3’ | qRT-PCR |
|  | TFAP2A reverse | 5’-AGGGGAGATCGGTCCGA-3’ | qRT-PCR |
| NR2F1 | NR2F1 forward | 5’-ATCGTGCTGTTCACGTCAGA-3’ | qRT-PCR |
|  | NR2F1 reverse | 5’-GCTCCTCACGTACTCCTCCA-3’ | qRT-PCR |
| SOX9 | SOX9 forward | 5’-GTACCCGCACTTGCACAAC-3’ | qRT-PCR |
|  | SOX9 reverse | 5’-TCTCGCTCTCGTTCAGAAGTC-3’ | qRT-PCR |
| chr1:9919431-9919620 | Chr1 forward | 5’-TGGGAAGGACACAGTGACAA-3’ | ChIP-qPCR |
|  | Chr1 reverse | 5’-CCTGGGTAAGGAGCTGGCTA-3’ | ChIP-qPCR |
| chr2:1484741-1484873 | Chr2 forward | 5’-ACTTCTGCTGTGGGCATGTT-3’ | ChIP-qPCR |
|  | Chr2 reverse | 5’-TGGGATCTCAAGCCAGCAAG-3’ | ChIP-qPCR |
| chr17:853129-853213 | Chr17 forward | 5’-CCTGGGCACCTTCTGGTAAC-3’ | ChIP-qPCR |
|  | Chr17 reverse | 5’-GGACCTGGGGCTAATTGCTT-3’ | ChIP-qPCR |
| chr18:37993305-37993425 | Chr18 forward | 5’-TTCAAGCAACAAGGCACAGG-3’ | ChIP-qPCR |
|  | Chr18 reverse | 5’-TTAGCTGCTGTGTGGCGTTA-3’ | ChIP-qPCR |
