## Supplementary figures and images for "Inability to switch from ARID1A-BAF to ARID1B-BAF impairs exit from pluripotency and commitment towards neural crest formation in *ARID1B*-related neurodevelopmental disorders"

### Supplementary_Fig_S1_resubmission.jpg

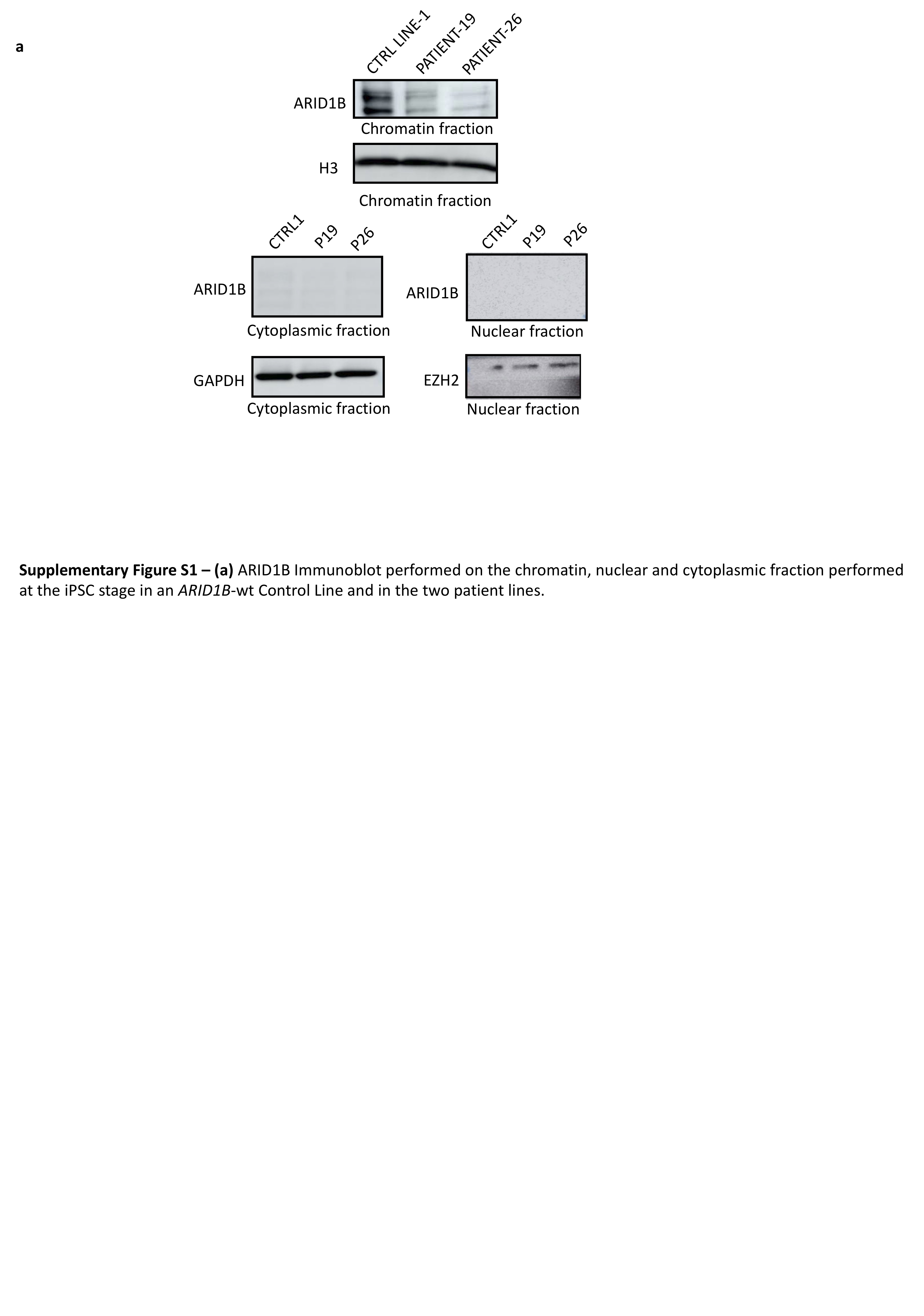

### Supplementary_Fig_S2.jpg

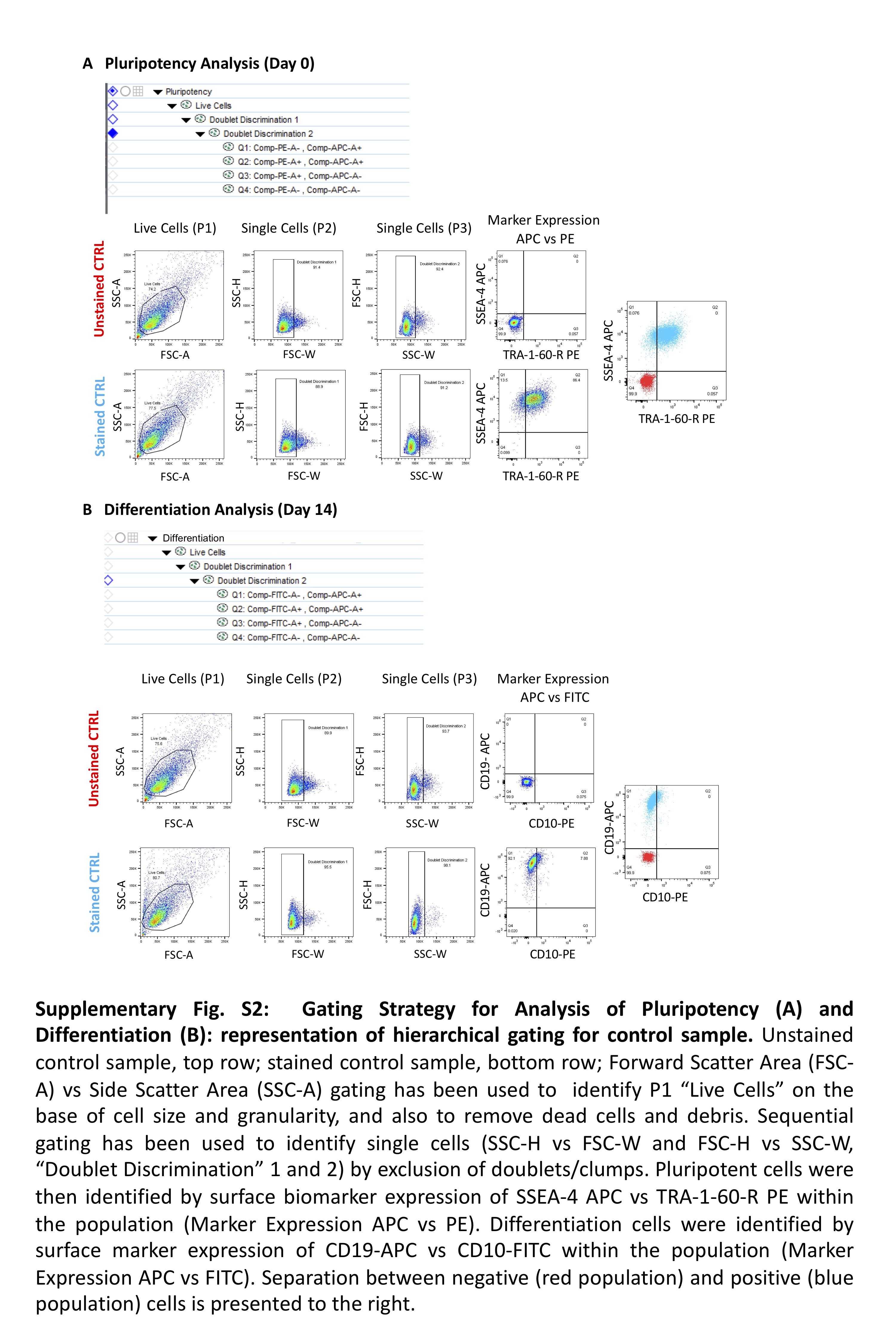

### Supplementary_Fig_S3_resubmission.jpg

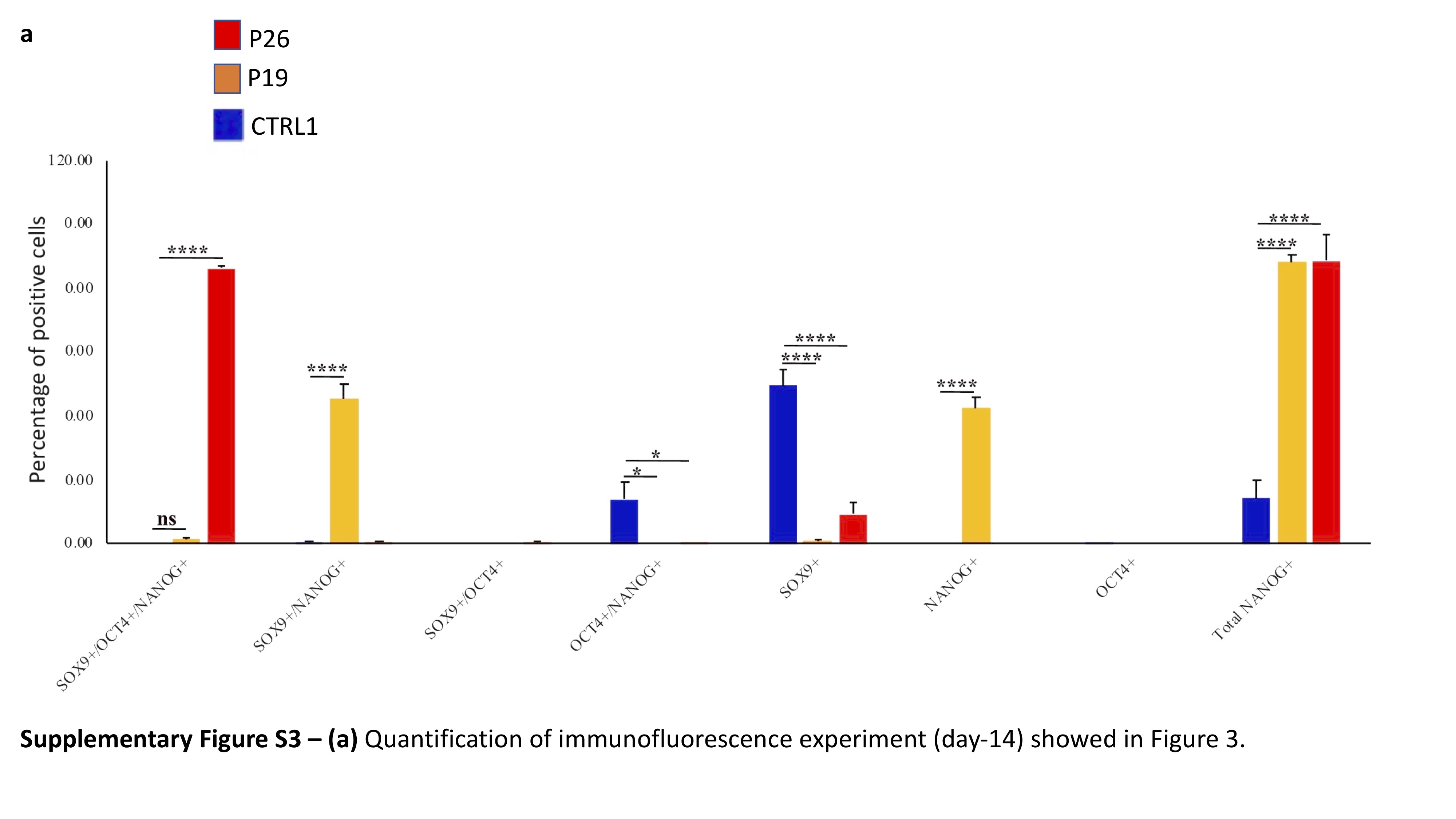

### Supplementary_Fig_S4_resubmision.jpg

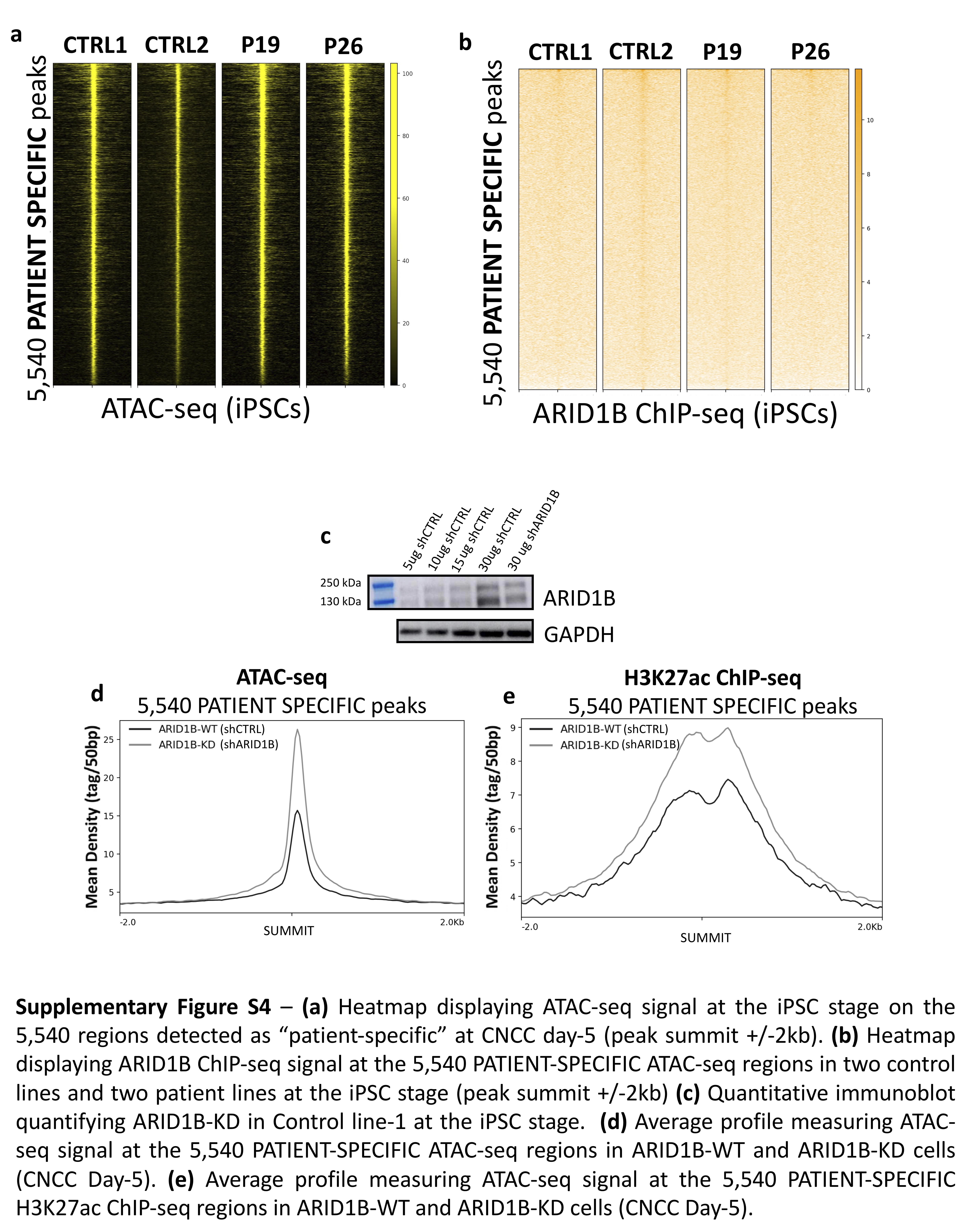

### Supplementary_Fig_S5.jpg

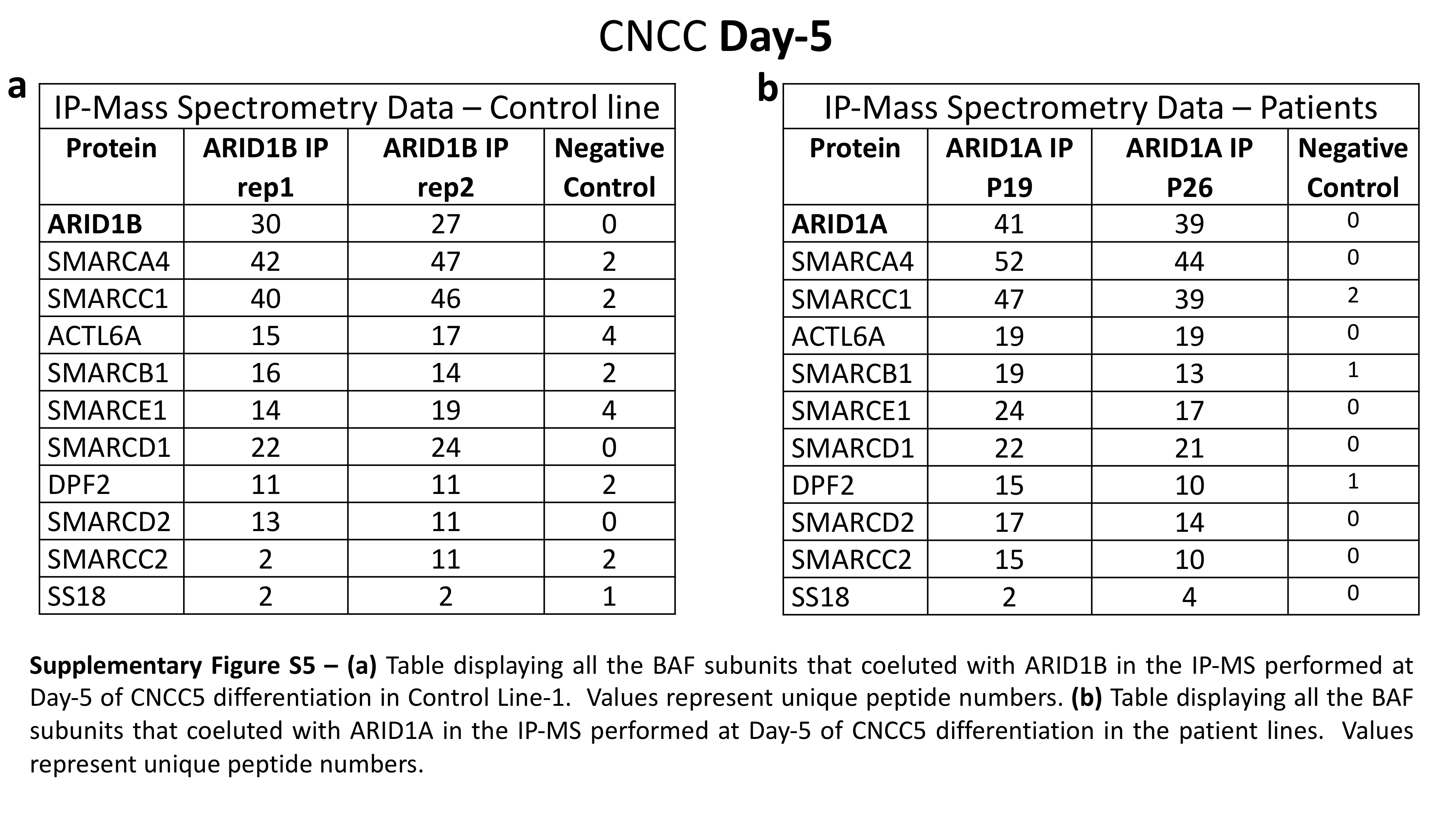
